# Optimization of Extraction Process by Response Surface Method-ology of Total Flavonoids from *Ziziphus Jujube* Flesh and its Sedative-Hypnotic Effects

**DOI:** 10.1101/2025.01.08.632005

**Authors:** Jie Li, Baojian Li, Xinbo Shi, Yuangui Yang, Zhongxing Song

## Abstract

**Background & aims:** This study optimizes the extraction process of total flavonoids from *Ziziphus jujube* flesh (TFZJF) by response surface methodology and investigating its sedative-hypnotic effects and mechanisms in a para-chlorophenylalanine (PCPA)-induced insomnia mouse model. It provides theoretical support for the further comprehensive utilization of *Ziziphus jujube* fruit and flesh.

**Methods:** Single-factor experiment and Box–Behnken response surface design were used to study the ethanol reflux extraction process of TFZJF, with flavonoid extraction rate as the indicator, to obtain the optimal extraction process of TFZJF. An insomnia model in mice was induced via an intraperitoneal injection of PCPA, and the effects of TFZJF on this model, along with its underlying mechanisms, were assessed using various approaches, including sodium pentobarbital-induced sleep potentiation, HE staining of tissue sections, ELISA, RT-PCR, WB, and serum metabolomics.

**Results:** The results showed that the optimized extraction conditions for TFZJF included a solid–liquid ratio (SLR) of 1:25 g·mL⁻1, ethanol concentration of 60%, and extraction time of 60 min, yielding an extraction efficiency of 1.98%. Data from the experimental groups indicated dose-dependent sleep improvement in the insomnia model, with the high-dose TFZJF (TFZJF-H) group exhibiting the most significant effect, followed by the medium-dose (TFZJF-M) and low-dose (TFZJF-L) groups. Metabolomic analysis revealed that TFZJF administration positively impacted the metabolic profile of PCPA-induced insomnia, particularly affecting pathways related to phenylalanine, tyrosine, cytochrome P450, and alanine metabolism on non-targeted metabolomics.

**Conclusions:** The extraction process is stable and reliable, which can be used for the extraction of TFZJF; meanwhile, TFZJF demonstrated significant sleep-enhancing effects in the PCPA-induced insomnia mouse model, supporting its potential for resource development and the utilization of non-medicinal parts of *Ziziphus jujube*.

## Introduction

*Ziziphus jujube* [*Ziziphus jujuba* Mill. var. *spinosa* (Bunge) Hu ex H. F. Chou], a woody plant of the Rhamnaceae family, mainly distributed as a food in Asia, Europe, and parts of the Americas. In China, it is mainly distributed in Shaanxi, Shanxi, Hebei, Shandong and other provinces, with an annual production exceeding 30,000 tons [1]. Due to its sour and sweet taste, jujube seed is widely eaten as a kind of fruit, and its seed jujube kernel is the preferred medicine for insomnia and widely used in the clinical prescription for insomnia. During the processing of jujube kernel, the pulp is discarded and not effectively utilized, resulting in waste of resources and damage to the ecological environment [2, 3]. Therefore, it is urgent to further study and effective utilization of sour jujube pulp. Ancient Chinese classics extensively document the medicinal properties of *Ziziphus jujube* seeds and flesh. In Shen Nong’s *Classic of the Materia Medica*, it is noted that *Ziziphus jujube* is “sour in taste and neutral in nature. Prolonged use supports the health of the five viscera, aids in weight reduction, and extends longevity,” marking the first recorded mention of its benefits. However, its application in treating insomnia was not included. In Zhang Zhong-jing’s Essentials from the Golden Cabinet (Han Dynasty), it was stated that “deficiency-consumption and vexation cause insomnia, and *Ziziphus Jujube* Decoction is used as treatment,” highlighting its therapeutic potential for insomnia, although the prescription itself does not specifically reference *Ziziphus jujube* seed. Further, the Newly Revised Materia Medica emphasizes the efficacy of *Ziziphus jujube* flesh in addressing insomnia without specifying seed usage, reflecting its historical role as a medicinal substance. Hence, *Ziziphus jujube* actually refers to the Ziziphus jujube flesh as recorded in Shen Nong’s *Classic of the Materia Medica*, indicating that the sour jujube pulp can also be used as medicine in history, and has related pharmacological activity.

Insomnia is a prevalent sleep disorder characterized by a range of physical, psychological, and behavioral clinical manifestations. Prolonged insomnia contributes to depression, exacerbates fatigue, impairs cognitive function, and compromises immune responses [4–6]. Contemporary studies highlight that neurotransmitter balance directly influences sleep quality [7,8]. Neurotransmitters, which mediate communication between neurons, are integral to brain function, particularly in sleep regulation. Their release and modulation are closely linked to sleep quality [9,10]. *Ziziphus jujube* seed, listed in the *Chinese Pharmacopoeia*, contains bioactive components such as flavonoids, saponins, alkaloids, and fatty acids, known for their sedative, antioxidant, anti-inflammatory, and cardioprotective properties [11]. The flesh of *Ziziphus jujube* also includes flavonoids, polysaccharides, alkaloids, and other compounds with sleep-enhancing, hepatoprotective, and anti-tumor activities [12]. Notably, flavonoids represent a key active component of *Ziziphus jujube* seed, and modern pharmacological research demonstrates their efficacy in improving sleep quality [13].

Currently, there is a paucity of research on the total flavonoids from *Ziziphus jujube* flesh (TFZJF), and no studies have reported the optimization of the extraction process for these compounds. The ethanol reflux extraction method is cost-effective in terms of equipment, features a straightforward operational workflow, exhibits excellent solubility, effectively separates impurities to enhance the purity of the target substance, and demonstrates both high extraction efficiency and yield for flavonoids [14]. Given that current studies typically use total flavonoid content as an extraction index, they often rely on single-factor experiments to assess how variables such as the extraction solvent, solid–liquid ratio (SLR), duration of extraction, and number of extractions influence total flavonoid recovery. A key advantage of response surface methodology lies in its ability to employ polynomial functions for fitting within the design space; this enables the precise approximation of functional relationships over localized areas with a minimal number of experiments while presenting results in a simple algebraic form. By selecting appropriate regression models, complex response relationships can be accurately fitted—demonstrating robust systematic properties that are practical and applicable across diverse fields [15]. *Ziziphus jujube* seed flavone enhances sleep quality by modulating brain levels of key neurotransmitters involved in sleep regulation [16,17]. Gamma-aminobutyric acid (GABA), an inhibitory neurotransmitter, diminishes neuronal excitability through its interaction with GABA receptors, inducing sedative and hypnotic effects. This is achieved by upregulating GABA_A_R1 expression, which directly influences GABA receptor availability [18–20]. Similarly, 5-hydroxytryptamine (5-HT), a neurotransmitter integral to the regulation of mood, sleep, and appetite, is widely recognized for its therapeutic potential in addressing insomnia, where increasing 5-HT levels in the brain is considered a beneficial approach [21–23]. Furthermore, research indicates that insomnia is linked to decreased levels of BDNF. Enhancing BDNF levels has been shown to improve sleep quality, underscoring BDNF’s role in supporting cognitive function and restoring healthy sleep patterns [24,25]. Flavonoids from *Ziziphus jujube* can penetrate the blood–brain barrier, directly impacting the central nervous system by modulating GABA, 5-HT, and other neurotransmitters, thus influencing sleep patterns. Moreover, they enhance sleep quality by affecting BDNF signaling pathways. These data suggest that *Ziziphus jujube* not only improves emotional stability but also supports neuronal survival and function, indicating its multifaceted role in neurotransmitter regulation to promote better sleep. Neurotransmitters and neurotrophic factors influenced by *Ziziphus jujube* are integral to the regulation of emotions, cognitive functions, and sleep [26–29]. Clinically, *Ziziphus jujube* seed has been widely used for treating various types of insomnia, with studies primarily focusing on the sedative and hypnotic mechanisms of its seed flavones. However, limited attention has been given to the sedative and hypnotic mechanisms of flavonoids found in *Ziziphus jujube* flesh.

Therefore, the present study utilized *Ziziphus jujube* waste material, specifically *Ziziphus jujube* flesh, as the experimental material and employed an ethanol reflux method for extraction. Based on the results of the single-factor experiment, the Box–Behnken response surface method was used to optimize the extraction process of TFZJF with flavonoid extraction rate as the indicator. Using *Ziziphus jujube* waste flesh as experimental materials, this study employed ethanol heating reflux for extraction. Following single-factor experiments, a Box–Behnken response surface methodology was applied to optimize the extraction conditions for TFZJF. The sedative and hypnotic effects of TFZJF were evaluated in insomnia models induced by intraperitoneal para-chlorophenylalanine (PCPA). Mice were allocated to a model group, diazepam (DZP) group, and TFZJF groups at low (TFZJF-L), medium (TFZJF-M), and high (TFZJF-H) doses. Sleep latency and duration were measured across all groups. Brain tissue samples were collected for histopathological examination via HE staining to observe morphological alterations. Elisa was employed to quantify 5-HT, GABA, and BDNF levels. Additionally, RT-qPCR and Western blot analyses were used to assess mRNA and protein expressions of 5-HT_1A_R, GABA_A_Rα1, and BDNF. Metabolomics and multivariate statistical methods were utilized to identify metabolic pathways and key metabolites in mouse serum following TFZJF treatment. This research provides a foundational understanding of the comprehensive use of non-medicinal parts of *Ziziphus jujube* based on the pharmacodynamics and mechanisms of TFZJF.

## Materials and Methods

### Materials

This study employed a variety of advanced instruments, including the 5810R high-speed refrigerated centrifuge (Eppendorf, Germany), the KQ-200TDE CNC ultrasonic cleaner (Kunshan Ultrasonic Instruments Co., Ltd. Jiangsu, China), and the 5430R high-speed centrifugal grinder (Eppendorf, Germany). Precision measurements were ensured using the Sar-torius CPA225D 1/100,000 electronic balance (Beijing Sartorius Scientific Instrument Co., Ltd. Beijing, China). Quantitative and qualitative analyses were performed with the Thermo Multiskan GO multifunctional microplate reader (Thermo Fisher Scientific, USA) and the qTOW-ER2.2 real-time fluorescence quantitative PCR instrument (ANALYTIK JEN, Germany), respectively. Microscopic observations were conducted using the Eclipse Ci-L upright white light camera microscope (Nikon, Japan), while absorption spectra were recorded via the UV-2600 ultraviolet–visible spectrophotometer (Shimadzu Corporation, Japan).

Rutin (purity ≥ 98%, HR1713S1, Baoji Herbest Bio-Tech Co., Ltd. Baoji, China), anhydrous ethanol, sodium nitrite, sodium hydroxide (analytical grade, Tianjin Tianli Chemical Reagent Co., Ltd. Tianjin, China), and aluminum nitrate (analytical grade, Chengdu Chrom Chemicals Co., Ltd. Chengdu, China) were utilized for experimental procedures. Additional reagents included PCPA (analytical grade, Sigma, USA), diazepam (2.5 mg/tablet, 220306, Shandong Xinyi Phar-maceutical Co., Ltd. Dezhou, China), and antibodies such as 5-HT_1A_R (AC231209084, Wuhan Servicebio Technology Co., Ltd. Wuhan, China), GABA_A_Rα1 (AC231216033, Wuhan Servicebio Technology Co., Ltd. Wuhan, China), and BDNF (AC231209053, Wuhan Servicebio Technology Co., Ltd. Wuhan, China). ELISA kits were sourced from Jiangsu Meimian Industrial Co., Ltd., including those for 5-HT (MM-0443M1, Jiangsu, China), GABA (MM-0442M1, Jiangsu, China), and BDNF (MM-0204M1, Jiangsu, China). RNA extraction from brain tissue was executed using the M5 HiPer Universal RNA Mini Kit (MF036, Mei5 Biotechnology, Beijing, China). For reverse transcription and quantitative PCR, the M5 Sprint qPCR RT kit with gDNA remover (MF166, Mei5 Biotechnology, Beijing, China) and the 2X M5 HiPer SYBR Premix EsTaq kit (MF787, Mei5 Biotechnology, Beijing, China) were employed. Staining was conducted using hematoxylin (CR2311076, Servicebio, Wuhan, China) and eosin (CR2402037-5, Servicebio, Wuhan, China).

The *Ziziphus jujube* sample, harvested in October 2022 from Hengshan District, Yulin, Shaanxi Province, was authenticated by Zhongxing Song, chief pharmacist at the Shaanxi Provincial- and Ministerial-Level Collaborative Innovation Center for Chinese Medicine Resources Industrialization. The specimen was confirmed as the dried fruit of *Ziziphus jujuba* Mill. var. *spinosa* (Bunge) Hu ex H.F. Chou, a member of the Rhamnaceae family.

### Assay of Total Flavonoid Content

#### Plotting of Standard Curve

The total flavonoid content was quantified using the NaNO2-Al (NO3)3-NaOH colorimetric method. A rutin reference solution (0.2 mg·mL^−1^) was prepared by dissolving the reference material in 65% ethanol. Specific volumes of the reference solution (0.0, 2.0, 3.0, 4.0, 5.0, 6.0, and 7.0 mL) were accurately transferred into 25 mL volumetric flasks. Subsequently, 1.0 mL of 5% NaNO2 solution was added, mixed thoroughly, and allowed to stand for 6 min. Next, 1.0 mL of 10% Al (NO3)3 solution was added, followed by another 6 min resting period after mixing. Finally, 10.0 mL of 4% NaOH solution was added, and the flasks were filled to the mark with distilled water. The mixtures were shaken well and left to stand for 15 min. Absorbance was then measured at 510 nm. A standard curve was plotted using rutin concentration on the X-axis and absorbance on the Y-axis, from which the regression equation was derived.

#### Determination of Total Flavonoid Content in Sample Solution

*Ziziphus jujube* flesh was ground, sieved through a 50-mesh sieve, and dried to a constant weight, yielding sample powder. A precise 1 g powder was weighed, and an appropriate solvent was added for heating reflux extraction under controlled conditions. The influence of varying ethanol concentrations, solid–liquid ratio (SLR), extraction times, and extraction frequencies on the flavonoid extraction from the flesh was systematically evaluated. The total flavonoid level was quantified with the linear equation derived from the standard curve, and the extraction efficiency was computed using the following formula:

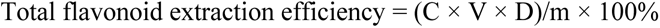

where C represented the total flavonoid concentration, mg·mL^−1^, determined from the standard curve; V denoted the extract volume, mL; D was the dilution factor; and m was the sample mass, mg.

### Preparation of Diazepam Gavage Solution

#### Single-Factor Experiment for Extraction Process

The influence of ethanol concentration (50%, 60%, 70%, 80%, and 90%), SLR (1:15, 1:20, 1:25, 1:30, and 1:35 mg·mL^−1)^, time of extraction (30, 60, 90, 120, and 150 min), and number of extraction cycles (1, 2, 3, 4, and 5) on the total flavonoid yield was evaluated. Each parameter was tested in triplicate for consistency.

#### Response Surface Optimization and Design of Extraction Process

Following the single-factor experiments, response surface design was applied using the experimental matrix outlined in Table 4. The extraction efficiency of total flavonoids (Y, %) was designated as the response variable. Verification experiments were subsequently performed based on the predicted optimal extraction parameters to determine the conditions that maximized flavonoid extraction efficiency.

### Drug Preparation

#### Preparation of PCPA Suspension

A 0.3% CMC-Na solution was prepared using physiological saline, heated in a water bath with magnetic stirring at 70 °C and 600 r·min^−1^. CMC-Na was gradually added until fully dissolved. Subsequently, 12 g PCPA powder was weighed and mixed with the CMC-Na solution and then diluted to a total volume of 300 mL. The suspension had a final concentration of (40 g·L^−1^).

#### Preparation of TFZJF Intragastric Solution

Ziziphus jujube flesh was dried, powdered, and subjected to ethanol reflux extraction under optimized conditions: 60% ethanol concentration, an SLR of 1:25 g·mL^−1^, and an extraction time of 60 min. The resulting extract was concentrated under reduced pressure to eliminate residual ethanol. Enrichment and purification were performed using D-101 macroporous resin. After loading 80 mL of the sample solution onto a preconditioned D-101 resin column, the pH was adjusted to 10. The sample was then eluted with 80% ethanol, and the eluate was concentrated and freeze-dried to yield TFZJF powder. The powder was dissolved in distilled water and stirred to prepare the TFZJF intragastric solution.

#### Preparation of Diazepam Intragastric Solution

Four diazepam tablets (10 mg per tablet) were pulverized into a fine powder, and purified water was added to reach a final volume of 50 mL, yielding a concentration of 0.1 mg·mL^−1^.

### Animals

A total of 60 SPF female KM mice (40 ± 2 g) were obtained from Chengdu Dossy Experimental Animals Co., Ltd, Chengdu, China. (Certificate number: SCXK (Chuan) 2020-0030). The mice were kept under controlled conditions at 23 ± 1.5 °C and 50–60% relative humidity, with no restriction to food or water, and maintained on a 12 h light/dark cycle. The experimental procedures were approved by the Ethics Committee of Shaanxi University of Chinese Medicine (approval number: SUCMDL20230913001).

### Animal Modeling and Administration

Female KM mice (n = 60) were randomly assigned to 6 groups, with 10 mice per group: (1) control, (2) model, (3) diazepam (DZP), (4) TFZJF-L, (5) TFZJF-M, and (6) TFZJF-H groups. The model was induced by intraperitoneal injection of PCPA at a dosage of 400 mg·kg^−1^ per day for four consecutive days [49]. During the modeling phase, all mice, except those in the control group, progressively exhibited behaviors such as mania, biting, increased aggression, disrupted circadian rhythms, dry fur, and decreased food consumption, confirming successful model establishment. Following this, drug administration was carried out via gavage once daily for 8 days. Mice in both the control and model groups received pure water, while the other groups received corresponding drugs: DZP at 1.3 mg·kg^−1^, TFZJF-L at 27 mg·kg^−1^, TFZJF-M at 54 mg·kg^−1^, and TFZJF-H at 81 mg·kg^−1^. After completing the sodium pentobarbital potentiation sleep experiment on the 8th day of treatment, the mice were euthanized via decapitation, and brain tissues were immediately collected. Some tissues were preserved in fixative for later use, while others were stored at −80 °C for later testing.

### HE Staining

Brain tissue was fixed for 48 h before embedding. Coronal sections, 3 μm in thickness, were prepared, dried at 45 °C for 1 h, and then subjected to dewaxing and dehydration. Hematoxylin staining was performed, followed by differentiation with ethanol. After rinsing, the sections were treated with 0.6% ammonia water for bluing, rinsed again, and subsequently stained with eosin. The samples were then dehydrated, cleared, and mounted with neutral gum. Morphological observations were conducted under a microscope at 200× magnification.

### ELISA Analysis

Brain tissue samples were homogenized, followed by centrifugation at 4 °C, 12,000 r·min^−1^ for 10 min (centrifugal radius: 9.5 cm). The supernatant was collected, and the ELISA procedure was conducted according to the kit’s protocol. The ELISA used in this experiment measures the content of 5-HT, GABA, and BDNF in the sample using a double antibody sandwich method. The capture antibodies for purified mouse 5-HT, GABA, and BDNF are coated onto microplates to form solid-phase antibodies. The microplates are then incubated with the sample, followed by the addition of HRP-labeled detection antibodies. An antibody-antigen-enzyme-labeled antibody complex is formed, which is then thoroughly washed. TMB is added to the wells, which turns blue in the presence of HRP enzyme and yellow in the presence of acid. The intensity of the color is directly proportional to the concentration of mouse 5-HT, GABA, and BDNF in the sample. The standard concentrations are 0, 0.5, 1, 2, 4, and 8 μmol/L. Each sample and standard are repeated three times. The absorbance at 450 nm (OD value) is measured using an enzyme-linked immunosorbent assay (ELISA) reader. The minimum detectable sensitivity of the 5-HT, GABA, and BDNF ELISA kit is less than 0.1 μmol/L. We used the standard concentrations and OD values to calculate the linear regression equation of the standard curve (R^2^ = 0.99). Then, the OD values of the samples were input into the equation to calculate the sample concentrations, which were multiplied by the dilution factor to obtain the actual concentrations. The levels of 5-HT, GABA, and BDNF in different animal groups were compared. Statistical analysis was performed using a t-test, with *p* < 0.05 indicating statistical significance.

### Western Blotting Analysis

Total protein from brain tissue was extracted by RIPA lysis buffer, and protein concentration was quantified with a BCA protein assay kit (Borst Biotechnology Co., Ltd., Wuhan, China). Proteins were resolved on 8–12% SDS-polyacrylamide gels and transferred onto PVDF membranes. Membranes were blocked with 5% skim milk for 2 h and then incubated overnight at 4 °C with primary antibodies: 5-HT_1A_R (1:500), GABA_A_Rα1 (1:800), and BDNF (1:500). After three washes, membranes were incubated with HRP-related secondary antibodies (1:1000 dilution) for 2 h. Visualization was performed using an enhanced chemiluminescence kit. Statistical analysis was conducted using t-test, with significance defined as *p* < 0.05.

### RT-qPCR Analysis

Total RNA extraction from brain tissue was carried out with the M5 HiPer Universal RNA Mini Kit (MF036, Mei5 Biotechnology, Beijing, China), followed by reverse transcription to cDNA utilizing the M5 Sprint qPCR RT Kit (MF166, Mei5 Biotechnology, Beijing, China) with gDNA removal. RT-qPCR was subsequently conducted with the 2× M5 HiPer SYBR Premix EsTaq Kit (MF787, Mei5 Biotechnology, Beijing, China). The 2^−△△Ct^ method (CT value comparison method) was used for relative quantitative analysis. The relative expression value obtained by inputting the average Ct value of the target gene (such as 5-HT_1A_R, GABA_A_Rα1, and BDNF) and the corresponding housekeeping gene (such as GAPDH) into the formula is the relative expression level. The formula is as follows: F = 2 − [[the average Ct value of the target gene in the test group − the average Ct value of the housekeeping gene in the test group] − [the average Ct value of the target gene in the control group − the average Ct value of the housekeeping gene in the control group]]. Each sample is repeated 3 times, and GAPDH, an internal reference gene, is added as a control to ensure the reliability and accuracy of the experimental results. The conditions for denaturation, annealing, and elongation during the PCR stage are as follows: 94 °C for 30 s, 55 °C for 30 s, and 72 °C for 30 s, for a total of 35 cycles. The primers for 5-HT1AR, GABAARα1, BDNF, and GAPDH were synthesized by Servicebio Co., Ltd. (Table 5.)

### Serum Metabolomics Analysis of TFZJF

#### Serum Sample Processing

For serum sample preparation, 50 µL of frozen serum was mixed with 150 µL of pre-cooled methanol and vortexed for 10 min. The mixture was then centrifuged at 10,000 r·min^−1^ for 10 min at 4 °C, with a centrifugal radius of 9.5 cm. The supernatant was achieved for further analysis.

#### Chromatography Mass Spectrometry Conditions

Mass spectrometry analysis and serum metabolite identification were conducted with UltiMate 3000 RS chromatograph coupled with the Q Exactive high-resolution mass spectrometer (Thermo Fisher Scientific, Shanghai, China). The chromatographic separation employed an AQ-C18 column (150 mm × 2.1 mm, 1.8 μm) with a mobile phase consisting of 0.1% formic acid (A) and methanol (B). A gradient elution profile was utilized: 5–55% B from 0 to 5 min, 55–65% B from 5 to 9 min, 65–85% B from 9 to 14 min, 85–90% B from 14 to 20 min, 90–95% B from 20 to 24 min, 95% B from 24 to 28 min, 95–5% B from 28 to 29 min, and 5% B from 29 to 30 min. Flow rate was 0.3 mL·min⁻¹, and volume was 5 μL, and the column temperature was set to 35 °C. Mass spectrometric detection was performed in both ion modes, with a scanning range of m/z 100–1500. High-purity nitrogen was supplied as the auxiliary spray ionization and desolvation gas, while nitrogen also served as the nebulizer and collision gas. Key parameters included a nebulizer temperature of 350 °C, nebulizer gas flow rate of 8 L·min⁻¹, sheath gas temperature of 350 °C, and sheath gas flow rate of 11 L·min^−1^. The capillary voltage was set to 3500 V for both ion modes, while nozzle voltages were adjusted to 1000 V for negative mode and 300 V for positive mode, with a pyrolysis voltage of 130 V.

#### Data Processing and Analysis

Chromatographic data from non-targeted metabolomics samples were acquired and processed using Xcalibur 4.1 (Thermo), with preliminary data sorting performed via CD3.3 (Thermo Fisher Scientific, Shanghai, China). Metabolite identification involved comparison with the mzCloud database. The processed data were further analyzed through PCA and OPLS-DA using SIMCA-P 14.1 software. Differential metabolites were screened by applying a VIP threshold of >1 and a significance level of p < 0.05. Potential biomarkers were subsequently confirmed through integration with the HMDB (https://hmdb.ca, accessed on 5 November 2023). Identified endogenous metabolites were mapped to relevant metabolic pathways via MetaboAnalyst 5.0, with impact > 0.1 as the criterion for pathway selection.

## Results

### Optimization of TFZJF Extraction Conditions by Single-Factor Experiment

The impact of ethanol concentration on the extraction of TFZJF is illustrated in Figure 1A. The extraction efficiency initially rose with an increasing ethanol concentration, peaking at 60%, before declining as the concentration continued to increase. Thus, 60% ethanol was selected for further experimentation. The effect of the SLR on extraction efficiency, with the optimal extraction occurring at a ratio of 1:25 g·mL^−1^ is shown in Figure 1B. Beyond this point, the efficiency decreased, confirming the ratio of 1:25 g·mL^−1^ as ideal for subsequent tests. The extraction time significantly influenced the extraction, with the maximum extraction achieved at 60 min, which was chosen for further analysis (Figure 1C). Increasing the number of extractions enhanced the extraction of TFZJF, stabilizing after two extractions (Figure 1D). Therefore, two extraction cycles were selected for the response surface experiment.

**Figure 1.**
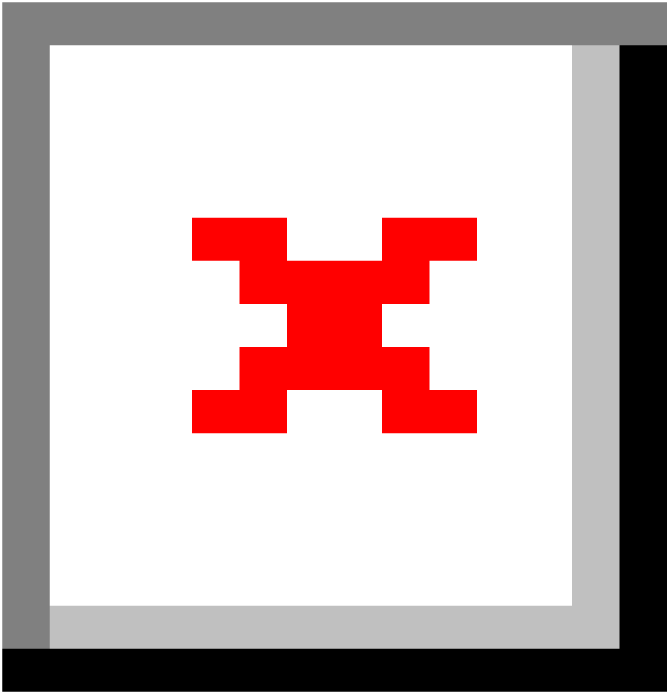
Single-factor investigation of TFZJF extraction conditions. (A) Single-factor investigation of ethanol concentration; (B) Single-factor investigation of solid–liquid ratio; (C) Single-factor investigation of extraction time; (D) Single-factor investigation of extraction times. The different colors represent the extraction rates of total flavonoids with different ethanol concentration, different solid–liquid ratio, different extraction time, and different extraction times.

### Statistical Analysis and Model Fitting

The response surface experimental design was utilized, with the results summarized in Table 1. Based on Design Expert 13, regression fitting analysis was applied, where the extraction time (A), ethanol concentration (B), and SLR (C) were considered the independent variables, and the total flavonoid extraction efficiency (Y) was the response variable. The resulting multivariate quadratic equation was expressed as follows: Y% = 1.92 − 0.0050A + 0.0325B + 0.0275C − 0.0075AB − 0.0875AC + 0.0175BC − 0.3488A2 − 0.1938B2 − 0.1387C2.

**Table 1.** Box–Behnken experimental design and results for TFZJF extraction conditions.

| Number | Extraction Time<br>(A, min) | Ethanol Concentration<br>(B, %) | Solid-to-Liquid Ratio<br>(C, g·mL<sup>-1</sup>) | Total Flavonoid Yield<br>(%) |
| --- | --- | --- | --- | --- |
| 1 | 60 | 70 | 20 | 1.58 |
| 2 | 60 | 60 | 25 | 1.89 |
| 3 | 90 | 60 | 30 | 1.37 |
| 4 | 60 | 70 | 30 | 1.67 |
| 5 | 90 | 60 | 20 | 1.49 |
| 6 | 60 | 60 | 25 | 1.97 |
| 7 | 60 | 50 | 20 | 1.54 |
| 8 | 60 | 60 | 25 | 1.91 |
| 9 | 30 | 60 | 30 | 1.55 |
| 10 | 30 | 70 | 25 | 1.42 |
| 11 | 90 | 50 | 25 | 1.35 |
| 12 | 60 | 60 | 25 | 1.95 |
| 13 | 60 | 60 | 25 | 1.88 |
| 14 | 30 | 60 | 20 | 1.32 |
| 15 | 90 | 70 | 25 | 1.39 |
| 16 | 60 | 50 | 30 | 1.56 |
| 17 | 30 | 50 | 25 | 1.35 |

The statistical analysis of the Analysis of Variance (ANOVA) is presented in Table 2. The p value of the model is less than 0.0001, indicating that the model has a significant effect on the extraction rate of TFZJF. The F and *p* values of the lack-of-fit term are 0.0556 and 0.9805, respectively, indicating that the lack-of-fit value is not significant. This confirms that the model has a good fit, with a small error, suggesting that the model is valid. The analysis of variance shows that the first-order terms B and C and the second-order terms A2, B2, and C2 all exhibit significant levels. In the regression model, R2 is 0.9929 and the adjusted R2adj is 0.9837, indicating that the model has a high goodness of fit and a high correlation between the predicted values and the measured values. The coefficient of variation (CV) is 1.87, indicating that the model has high precision and reliability. In conclusion, the established model of the extraction rate of TFZJF has high accuracy and credibility and can be used for predictive analysis of the extraction amount of TFZJF. The three first-order terms (A, B, C), the three second-order terms (AB, AC, BC), and A2, B2, and C2 all have a relevant influence on the extraction rate of TFZJF, indicating that the influence of the three factors on the extraction rate of TFZJF is not a simple linear function relationship. The F values of the three factors are B > C > A, indicating that the order of the influence of each factor on the extraction rate of TFZJF is ethanol concentration > SLR > time. The extraction time and ethanol concentration, SLR (AB, AC), and ethanol concentration and SLR (BC) also have a relevant influence on the extraction rate of TFZJF.

**Table 2.** Analysis of variance results.

|  | Sum of Squares | Mean Square | *F* Value | *p* Value |
| --- | --- | --- | --- | --- |
| Model | 0.8696 | 0.0966 | 108.22 | <0.0001 |
| A | 0.0002 | 0.0002 | 0.224 | 0.6504 |
| B | 0.0084 | 0.0084 | 9.46 | 0.0179 |
| C | 0.006 | 0.006 | 6.78 | 0.0353 |
| AB | 0.0002 | 0.0002 | 0.252 | 0.6311 |
| AC | 0.0306 | 0.0306 | 34.3 | 0.0006 |
| BC | 0.0012 | 0.0012 | 1.37 | 0.2798 |
| A<sup>2</sup> | 0.5121 | 0.5121 | 573.57 | <0.0001 |
| B<sup>2</sup> | 0.1581 | 0.1581 | 177.03 | <0.0001 |
| C<sup>2</sup> | 0.0811 | 0.0811 | 90.79 | <0.0001 |
| Residual error | 0.0063 | 0.0009 |  |  |
| Lack of fit | 0.0003 | 0.0001 | 0.0556 | 0.9805 |
| Pure error | 0.006 | 0.0015 |  |  |
| R<sup>2</sup> | 0.9929 |  |  |  |
| R<sup>2</sup><sub>adj</sub> | 0.9837 |  |  |  |
| CV | 1.87 |  |  |  |

### Response Surface Analysis of TFZJF Extraction Process

To enhance the understanding of the interactions among test variables, response surface and contour plots (Figure 2) were generated to illustrate the effects of the extraction time, ethanol concentration, and SLR on the extraction rate of TFZJF. The response surfaces for these interactions exhibited a downward opening, indicating that the extraction rate of TFZJF initially increased before subsequently decreasing as A, B, and C were elevated. The interaction effects of the ethanol concentration and extraction time on the TFZJF extraction are depicted in Figure 2A, B. The flavonoid extraction rate increased markedly as the ethanol concentration approached 60% and the extraction time reached 60 min, after which a decline was observed. The influence of the SLR and extraction time at a fixed ethanol concentration of 60% is illustrated in Figure 2C, D, with the highest extraction efficiency achieved at an SLR of 1:25 g·mL^−1^ and an extraction time of 60 min. Similarly, the combined effects of the SLR and ethanol concentration on flavonoid extraction at an extraction duration of 60 min is demonstrated in Figure 2E, F, indicating that the optimal extraction efficiency occurred when the SLR was 1:25 g·mL^−1^ and ethanol concentration was 60%. The response surface model predicts the optimal extraction process as follows: a 59.334 min extraction time, 60.87% ethanol concentration, SLR of 25.562:1 g·mL^−1^, and total flavonoid extraction rate of 1.923%. Considering operational errors, the optimized experimental conditions are as follows: a 60 min extraction time, 60% ethanol concentration, and SLR of 25:1 g·mL^−1^. Three verification experiments were conducted under these conditions, and the calculated total flavonoid extraction rate was 1.983 ± 0.012%, with a small error from the predicted value, further verifying the reliability and stability of the model.

**Figure 2.**
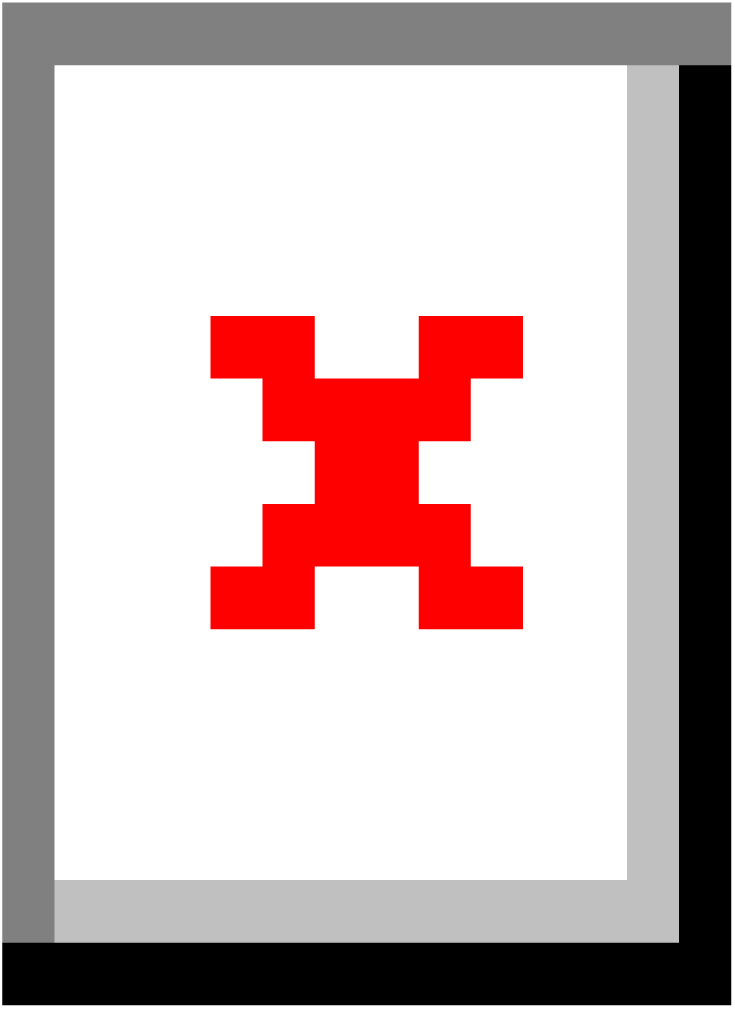
Results of the interaction of various extraction factors of TFZJF. (A) Response surface diagram of ethanol concentration and extraction time to total flavonoids extraction rate; (B) Ethanol concentration and extraction time on total flavonoids extraction rate contour map; (C) Solid–liquid ratio and extraction time to total flavonoids extraction rate response surface; (D) Solid–liquid ratio and extraction time to total flavonoids extraction rate contour map; (E) Solid–liquid ratio and ethanol concentration on the response surface of total flavonoids extraction rate; (F) Solid–liquid ratio and ethanol concentration on total flavonoids extraction rate contour map. Note: The red dot in the center of the graph represents the optimal condition where two conditions intersect, and the red dots around it represent the range of the interaction conditions.

### Results of TFZJF Purification Process

#### Adsorption and Desorption Rates of Various Macroporous Resins

Five types of macroporous resins—HPD-600, HPD-100, D-101, HP-20, and AB-8—were evaluated based on their adsorption and desorption rates. These parameters served as the criteria for selecting the optimal resin model. The analysis revealed that D-101 resin exhibited superior adsorption and desorption rates, leading to its selection as the most suitable resin for the adsorption process (Figure 3).

**Figure 3.**
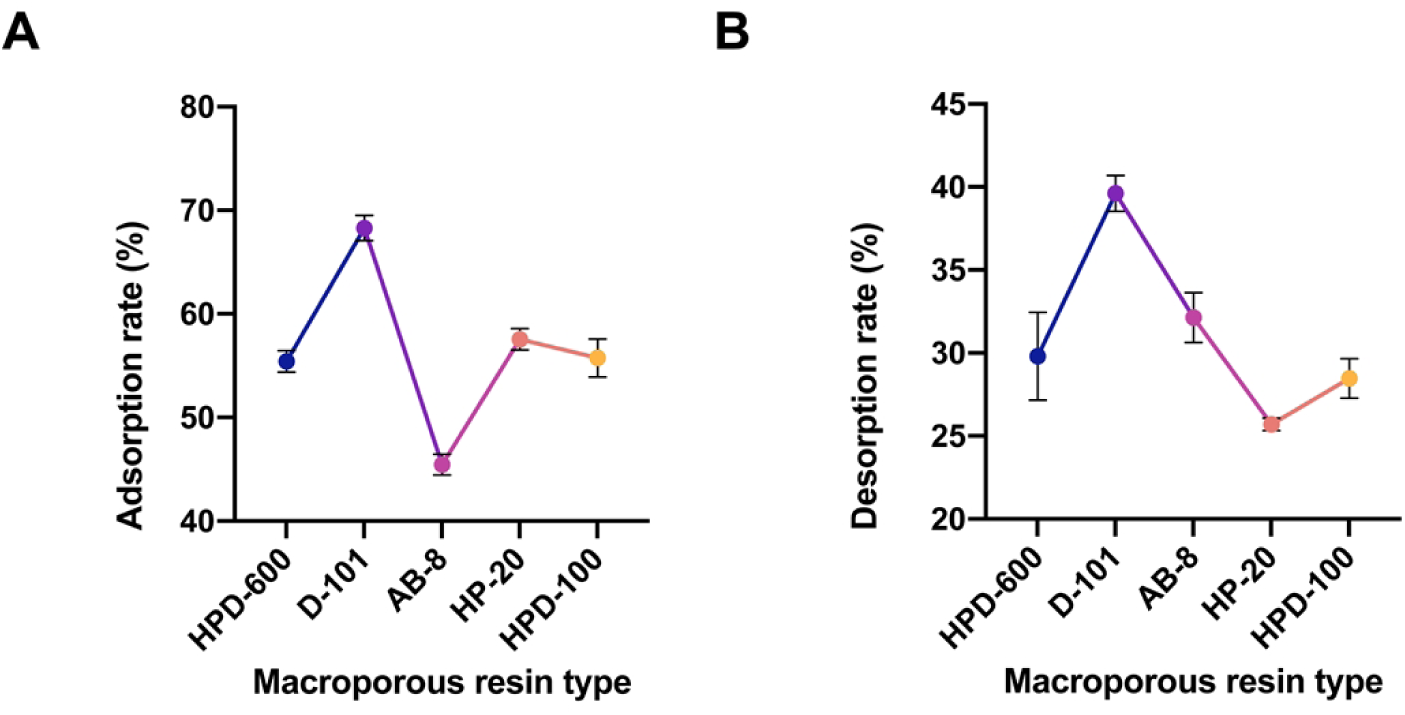
Screening of macroporous resin type. (A) Adsorption rate; (B) Desorption rate. Note: Blue represents the adsorption rate and desorption of the HPD-600 type; purple represents the adsorption rate and desorption of the D-101 type; pink represents the adsorption rate and desorption of the AB-8 type; orange represents the adsorption rate and desorption of the HP-20 type; yellow represents the adsorption rate and desorption of the HPD-100 type.

#### Effect of *Ziziphus jujube* Flesh Extract on Adsorption Rate at Different pH Values

Figure 4A displays the adsorption rate of *Ziziphus jujube* flesh extract at various pH levels. The adsorption rate was notably higher in alkaline conditions, peaking at pH 10, which was subsequently chosen for macroporous resin elution. As depicted in Figure 4B, the maximum adsorption capacity was reached when 40 mL of sample with a 10% concentration approached the theoretical leakage point. At 80 mL, the resin reached its saturation point, and further addition of *Ziziphus jujube* extract resulted in an incomplete adsorption. Figure 4C shows the desorption rate across ethanol concentrations, where a gradual increase was observed as ethanol concentration rose from 20% to 80%. The desorption rate peaked at 80% ethanol. Following the optimal extraction process, the extract was concentrated under reduced pressure to remove residual alcohol. The pH was adjusted to 10 before being adsorbed onto a pretreated D-101 macroporous resin column, eluted with 80% ethanol, and the eluate was collected and concentrated. Total flavonoid content analysis revealed that the purity of TFZJF purified by D-101 macroporous resin reached 58.32%.

**Figure 4.**
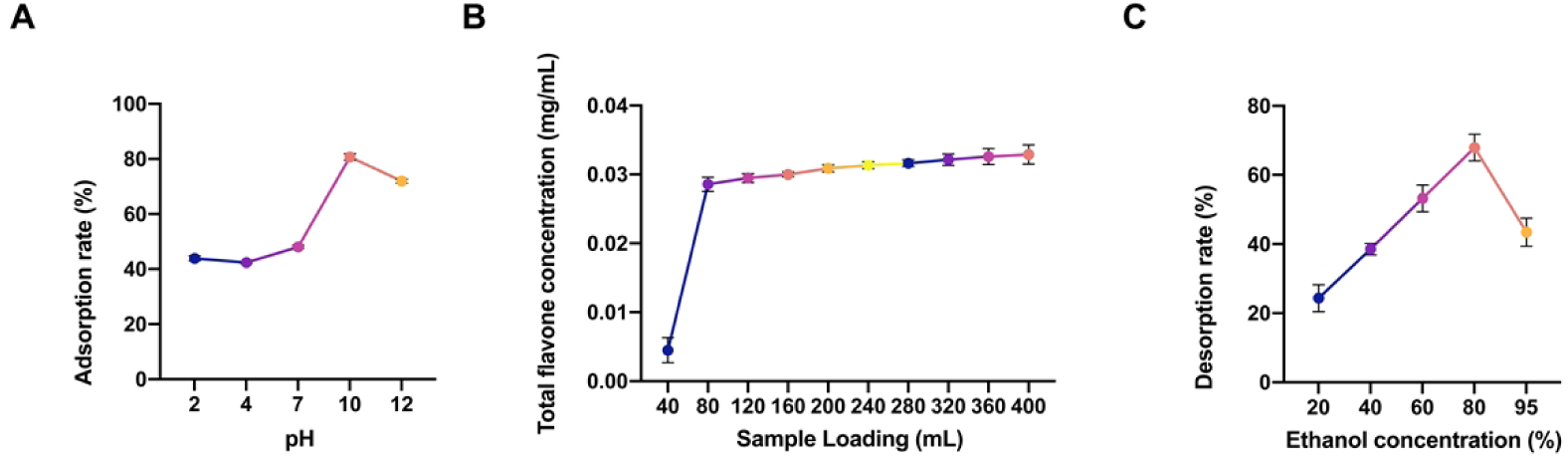
Results from the investigation of the TFZJF purification process. (A) Adsorption rate under different pH conditions; (B) Maximum adsorption capacity; (C) Desorption rates of different ethanol concentrations.

### Sleep Latency and Sleep Duration After TFZJF Treatment

Out of 60 mice, 10 were randomly assigned to the blank group, while the remaining 50 were allocated to the model group, DZP group, TFZJF-L, TFZJF-M, and TFZJF-H groups following successful induction of an insomnia model via an intraperitoneal injection of PCPA. Continuous gastric administration was applied to all groups. After 8 days of treatment, a sodium pentobarbital potentiation sleep experiment was conducted, and sleep latency and duration were recorded for each group. As presented in Figure 5, the model group exhibited a significant increase in sleep latency compared to the blank group (*p* < 0.01). Sleep latency in the DZP and TFZJF dosage groups was significantly reduced compared to the model group (*p* < 0.01), with TFZJF-H showing the shortest latency among the TFZJF dosage groups, followed by TFZJF-M, with TFZJF-L displaying the longest. Figure 5 showed that the model group experienced a marked reduction in sleep duration relative to the blank group (*p* < 0.01). Sleep duration in the DZP, TFZJF-H, and TFZJF-M groups significantly increased compared to the model group (*p* < 0.01), while the increase in the TFZJF-L group was not statistically significant. Among the TFZJF dosage groups, TFZJF-H resulted in the longest sleep duration, followed by TFZJF-M, with TFZJF-L having the shortest.

**Figure 5.**
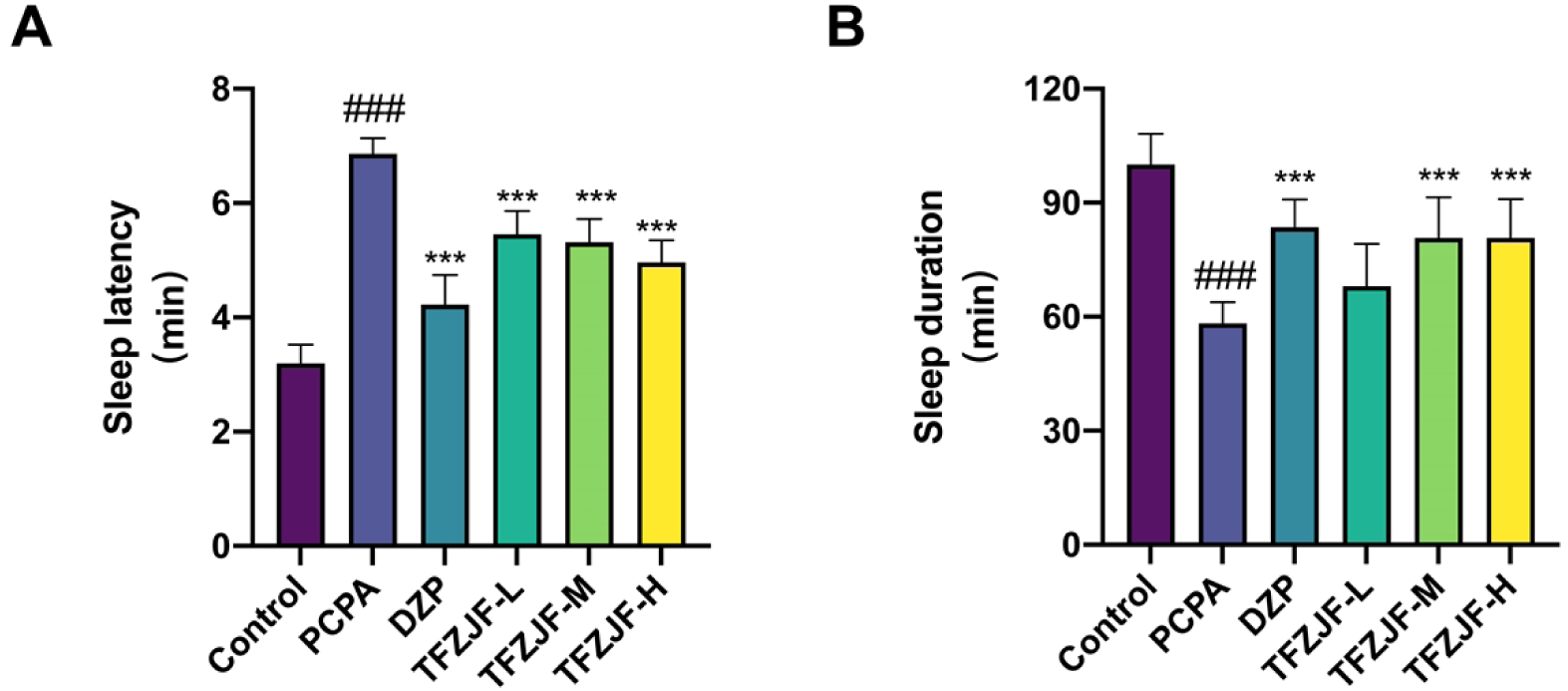
Effects of TFZJF on (A) sleep latency and (B) sleep duration in PCPA-induced insomnia model mice. Data were presented as mean ± SD. Comparisons to the control group show ^###^ *p*< 0.001; comparisons to the PCPA group indicate *** *p*<0.001.

### HE Staining Analysis

As depicted in Figure 6, HE staining revealed irregular cellular arrangement in the brain tissue of the model group compared to the blank group. In the hypothalamus, numerous neurons exhibited shrunken, intensely stained nuclei, deformed cell bodies, and indistinct boundaries between the nucleus and cytoplasm. Additionally, extensive neuronal degeneration was observed, characterized by loose, lightly stained cytoplasm. In contrast, brain tissue from the DZP and TFZJF treatment groups displayed clearer staining. Neuronal nuclei shrinkage was partially alleviated in the hypothalamus, with reduced signs of degeneration. These results indicated that TFZJF contributed to the improvement of brain tissue morphology in mice.

**Figure 6.**
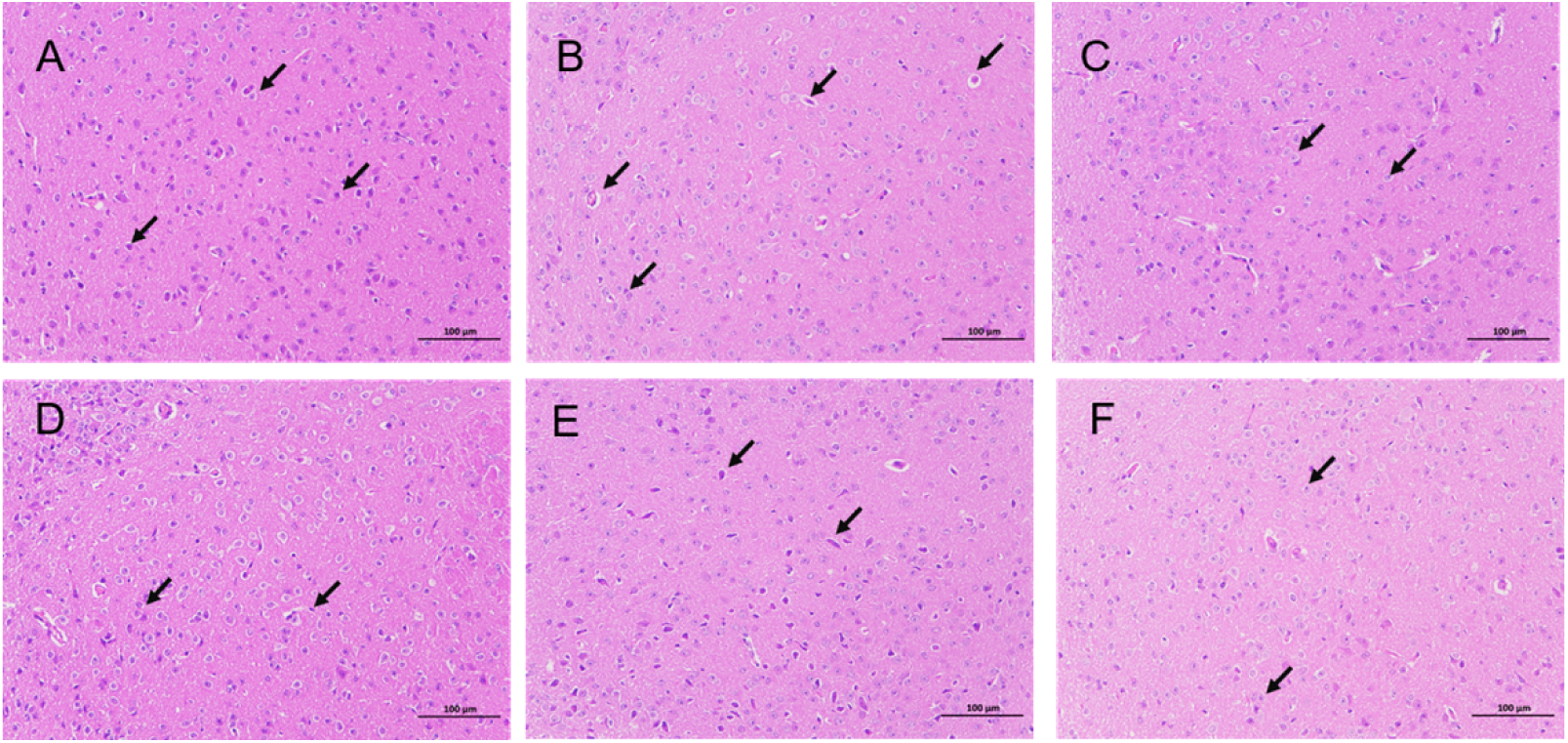
Effect of TFZJF on pathological morphology of mouse brain tissue (HE, ×200). (A) Control; (B) model; (C) DZP; (D) TFZJF-H; (E) TFZJF-M; (F) TFZJF-L. Note: The black arrows indicate the pathological changes in the nucleus of the hypothalamus neurons in the brain tissue.

### ELISA Results

The ELISA analysis, as shown in Figure 7, revealed significant reductions in 5-HT, GABA, and BDNF levels in the model group compared to the blank group (*p* < 0.01). However, the administration of DZP, TFZJF-H, and TFZJF-M resulted in significant increases in 5-HT, GABA, and BDNF levels (*p* < 0.01) compared to the model group. In contrast, the TFZJF-L group did not show a statistically significant increase in these biomarkers. Among the TFZJF dosage groups, the 5-HT, GABA, and BDNF contents followed a descending order: TFZJF-H, TFZJF-M, and TFZJF-

**Figure 7.**
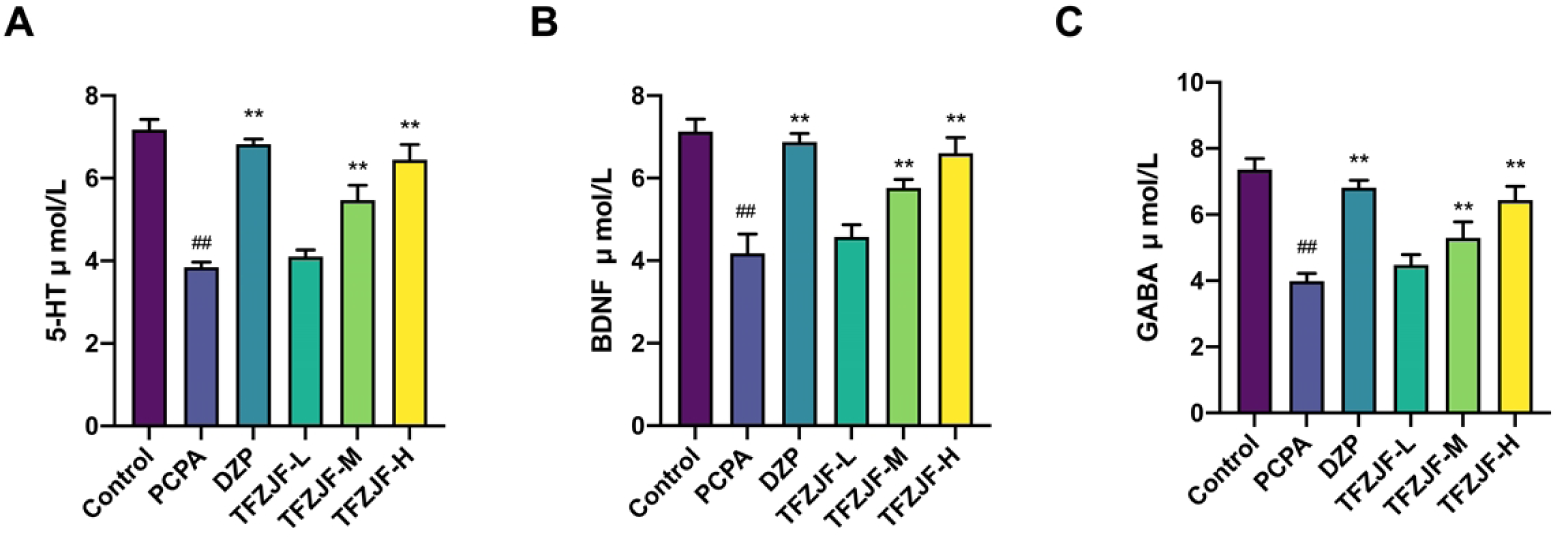
Measurement of (A) 5-HT, (B) BDNF, and (C) GABA content in the brain tissue of mice in each group by ELISA. Data were presented as mean ± SD. Comparisons to the control group show, ^##^ *p* < 0.01; comparisons to the PCPA group indicate, ** *p* < 0.01, n = 6.

### PCR Analysis

The analysis of the PCR results, as presented in Figure 8, revealed a significant reduction in the mRNA expression levels of 5-HT_1A_R, GABA_A_Rα1, and BDNF in the model group compared to the blank group (*p* < 0.01). In contrast, mRNA expression levels of 5-HT_1A_R, GABA_A_Rα1, and BDNF in the DZP, TFZJF-H, and TFZJF-M groups were notably increased relative to the model group (*p* < 0.05). The increase in mRNA expression in the TFZJF-L group, however, was not statistically significant. Among the different TFZJF dosage groups, the expression levels of 5-HT_1A_R, GABA_A_Rα1, and BDNF followed a descending order: TFZJF-H, TFZJF-M, and TFZJF-L.

**Figure 8.**
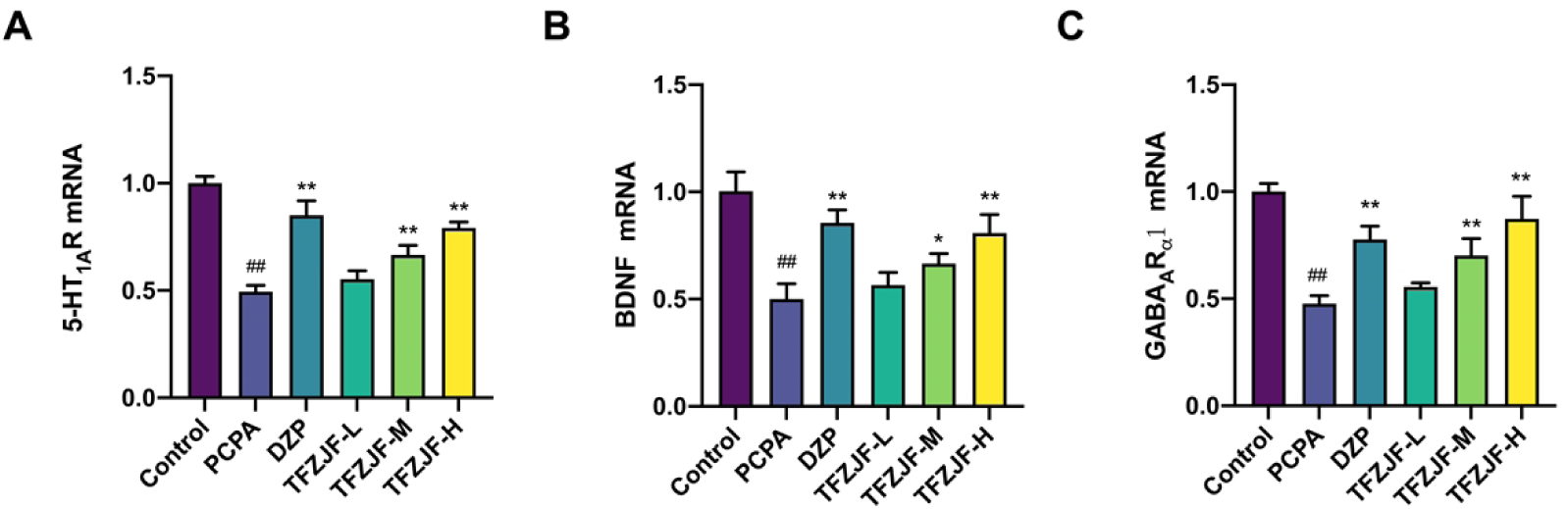
The mRNA expression levels of 5-HT_1A_R, BDNF, and GABA_A_Rα1 in each group of mice measured by PCR. (A) The mRNA expression level of 5-HT_1A_R; (B) The mRNA expression level of BDNF; (C) The mRNA expression level of GABA_A_Rα1. Data were presented as mean ± SD. Comparisons to the control group show, ^##^ *p* < 0.01; comparisons to the PCPA group indicate * *p* < 0.05, ** *p* < 0.01, n = 3.

### Western Blotting Results

As depicted in Figure 9, WB analysis revealed that the model group exhibited a significant reduction in the protein expression of 5-HT_1A_R, GABA_A_Rα1, and BDNF compared to the blank group (*p* < 0.01). The treatment with DZP, TFZJF-H, and TFZJF-M significantly upregulated the expression of 5-HT_1A_R, BDNF, and GABA_A_Rα1 relative to the model group (*p* < 0.05). These results suggested that both medium and high doses of TFZJF exerted anti-insomnia effects, potentially through the modulation of 5-HT_1A_R, GABA_A_Rα1, and BDNF expression.

**Figure 9.**
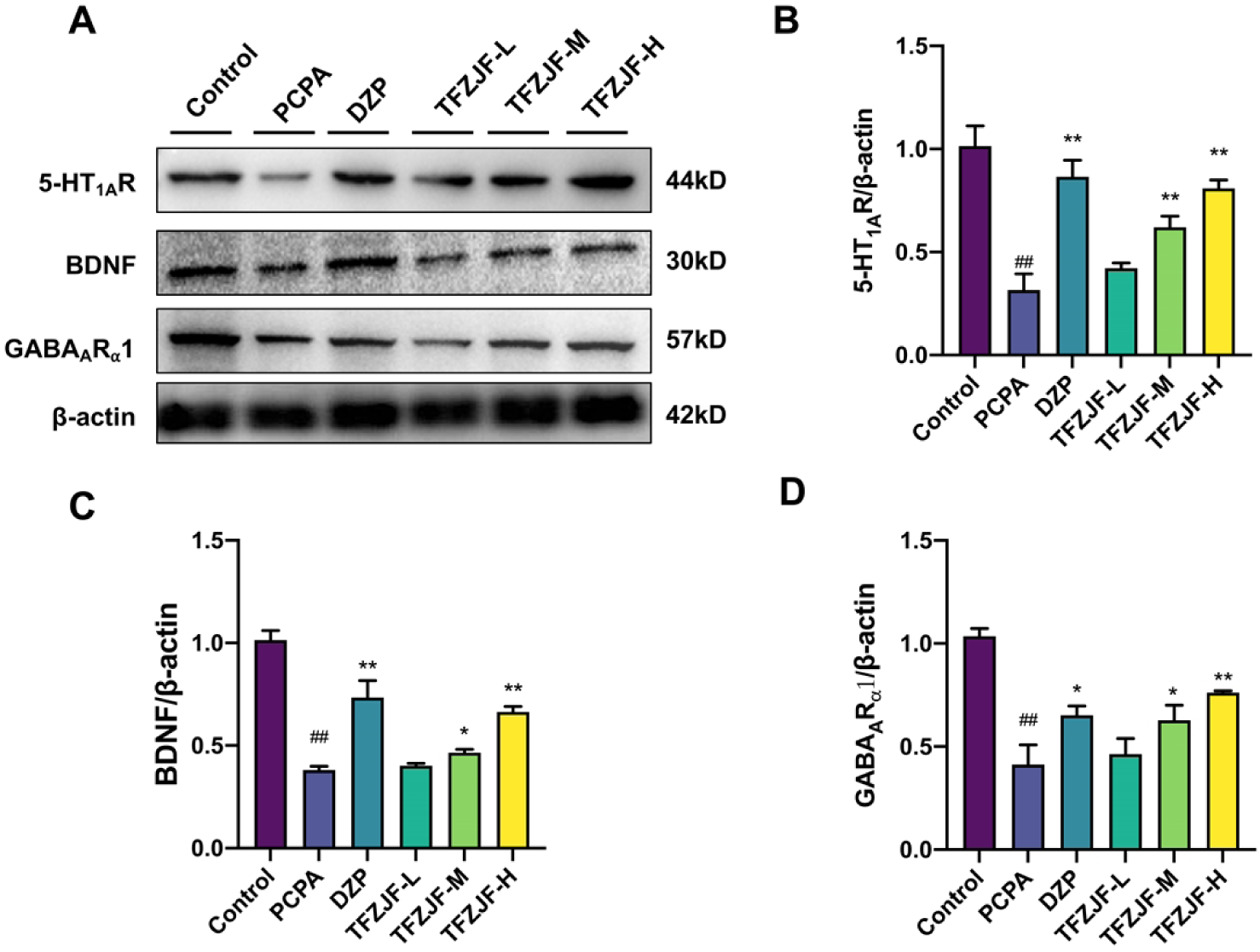
(A) Western blotting analysis of 5-HT_1A_R, BDNF, and GABA_A_Rα1 in mice of each group; (B) proportion of immunoblotting bands of 5-HT_1A_R, (C) BDNF to Beta relative to Beta Actin and (D) GABA_A_Rα1. Data were presented as mean ± SD. Comparisons to the control group show, ^##^ *p* < 0.01; comparisons to the PCPA group indicate * *p* < 0.05, ** *p* < 0.01, n = 3.

### TFZJF Serum Metabolomics Analysis

Metabolomics and multivariate statistical analysis, based on UPLC-Q-TOF-MS, were employed to assess alterations in serum metabolomics of mice following the administration of TFZJF. PCA of the non-targeted metabolomics profile revealed significant distinctions between the control and model groups (Figure 10A), confirming the successful establishment of the PCPA-induced insomnia mouse model. Comparative analysis across the control, model, and treatment groups, using PCA (Figure 10B) and cluster heat map analysis (Figure 10D), demonstrated tight clustering within each group, indicating minimal intragroup variability. Furthermore, the proximity of metabolite clusters in the treatment group to those in the control group suggested a normalization of the metabolic profile in PCPA-induced insomnia mice following administration. PLS analysis was used to generate VIP scores for each metabolite, identifying those most relevant to the separation of the two groups in both negative and positive ion modes (Figure 10C). To further elucidate the metabolic pathways affected by TFZJF, MetaboAnalyst 6.0 was applied to analyze significantly altered metabolites. Differentially expressed metabolites (DEMs) between the control and model groups, along with those reversed by flavonoid treatment, were detailed in Table 3. The non-targeted metabolomics results (Figure 10E) indicated that the most affected pathways included phenylalanine, tyrosine, cytochrome P450, and alanine metabolisms.

**Figure 10.**
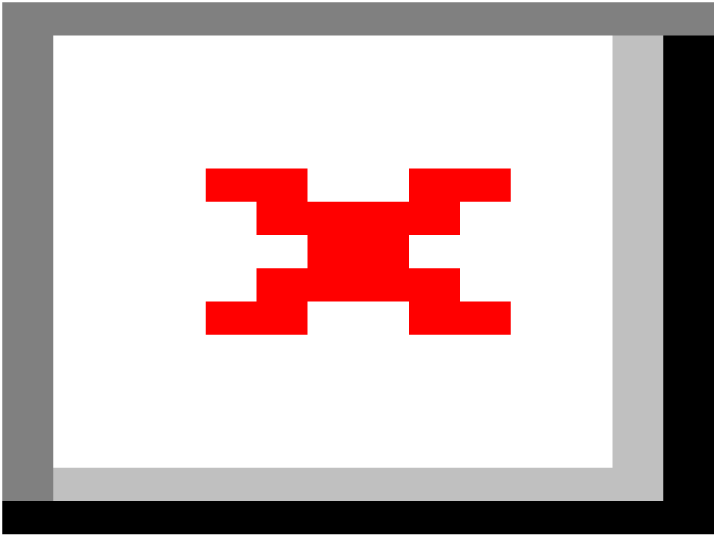
Effect of TFZJF on the differential metabolite expression in PCPA−induced insomnia model mice. (A) PCA plot of the DEMs in each group; (B) PLS−DA in each group; (C) VIP results; (D) heatmap of the DEMs in each group; (E) metabolic pathway enrichment analysis diagram of the DEMs. Note: In the metabolic pathway enrichment bubble plot, the color of the bubbles represents the significance of enrichment: the color of the bubbles reflects the size of the *P*−value, and the redder the color, the larger −log10 (*P*−value), and the whiter the color, the smaller −log10 (*P*−value).

**Table 3.**
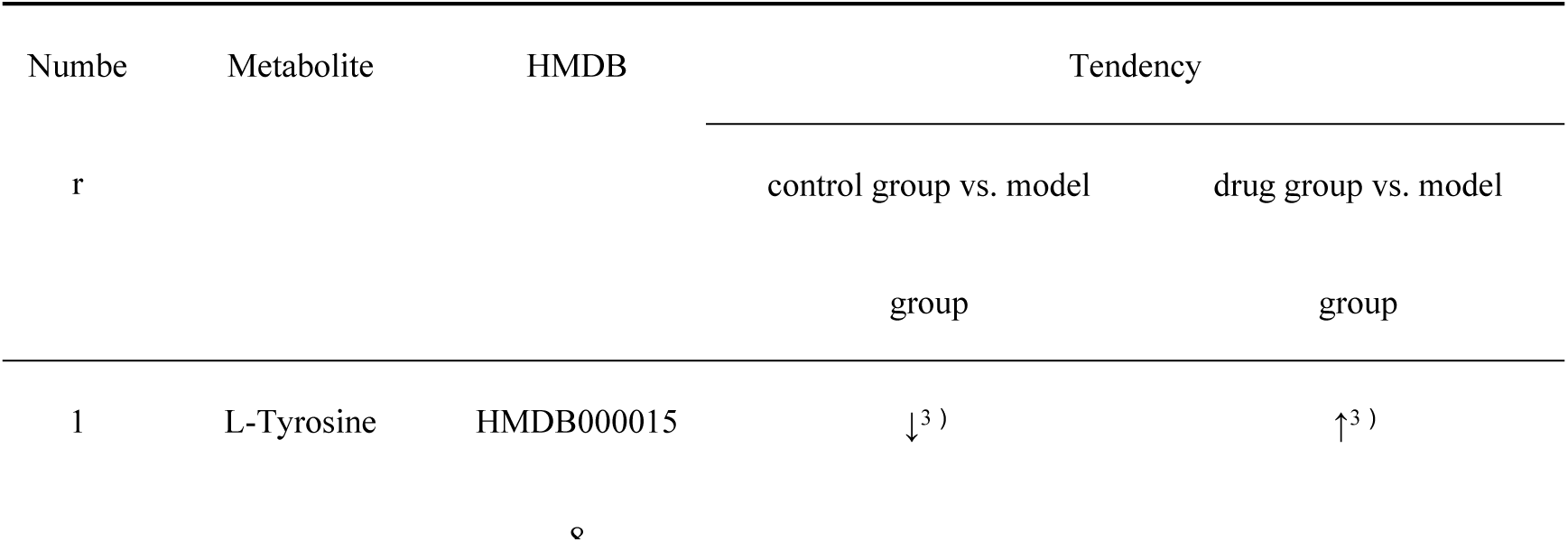
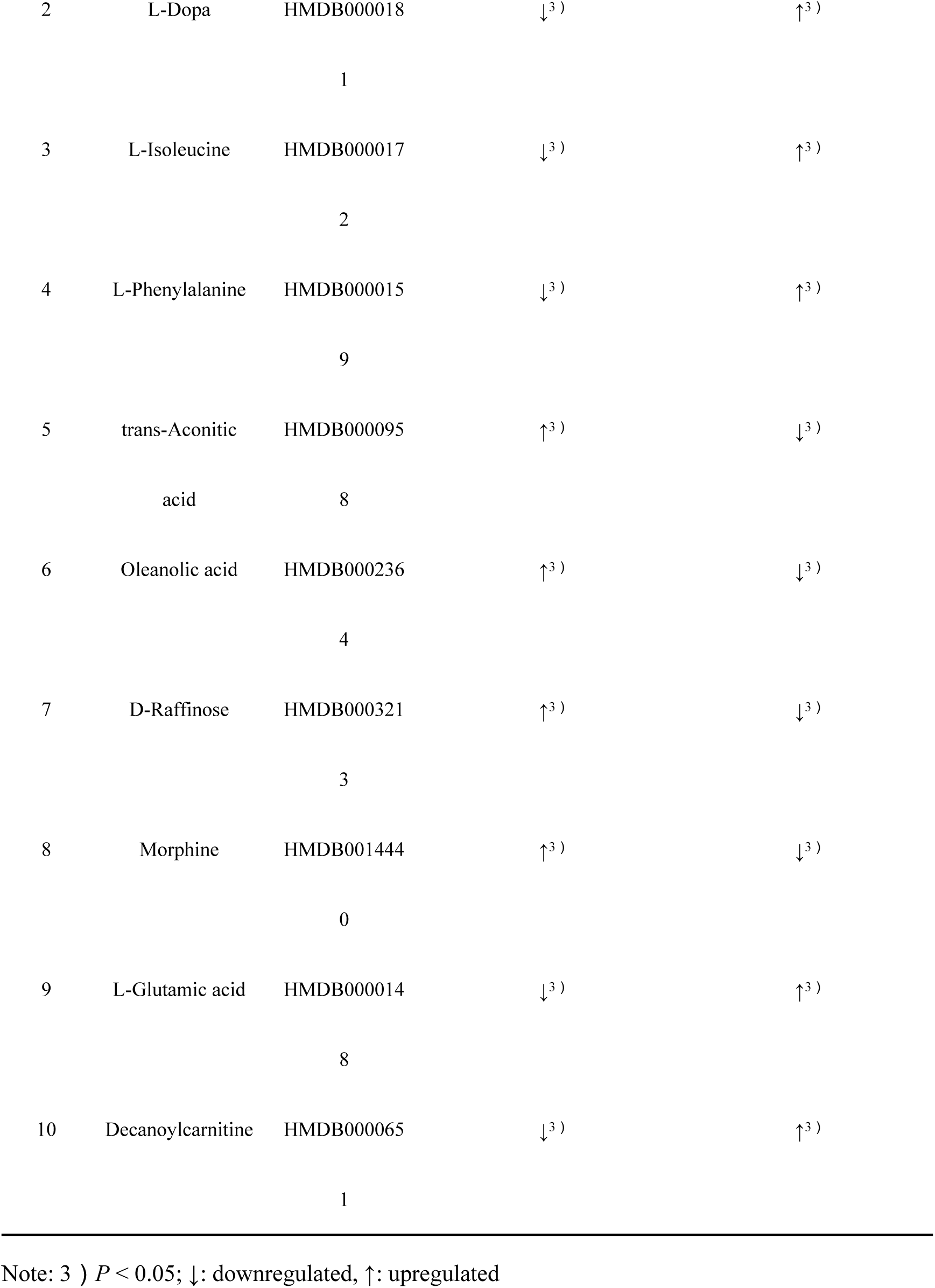
Comparison of DEM in control; Model and TFZJF administration group.

**Table 4.** Factors and levels for response surface design.

| Level | Factor |  |  |
| --- | --- | --- | --- |
|  | Extraction Time | Ethanol Concentration | Solid-to-Liquid Ratio |
|  | (A, min) | (B, %) | (C, g·mL<sup>-1</sup>) |
| -1 | 30 | 30 | 1:20 |
| 0 | 60 | 60 | 1:25 |
| 1 | 90 | 90 | 1:30 |

**Table 5.**
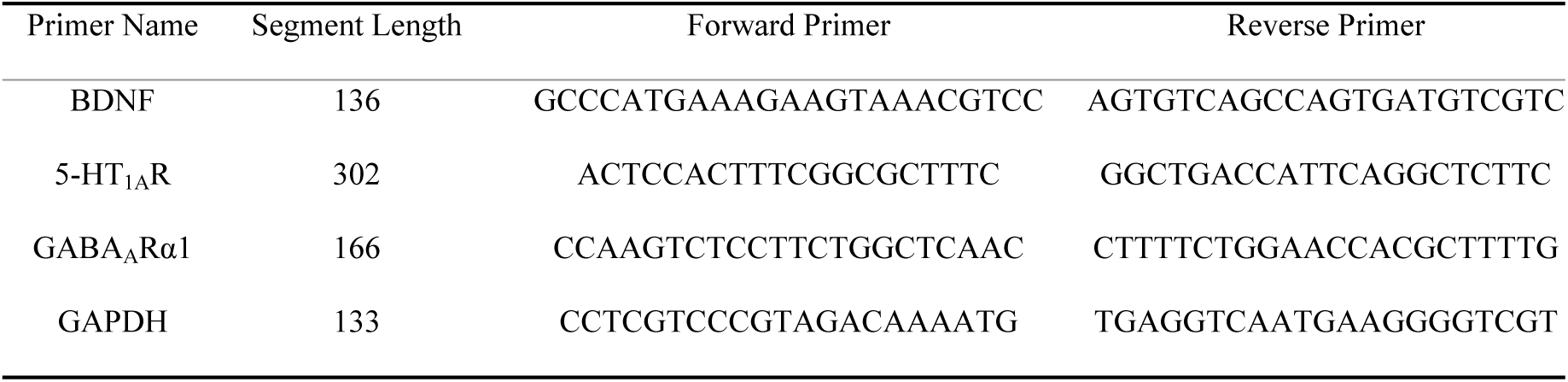
Primers sequence.

## Discussion

Flavonoids present in Ziziphus jujube seed are known for their sedative and hypnotic effects. While previous studies have confirmed the presence of flavonoids in Ziziphus jujube flesh, research on TFZJF remains limited. Ziziphus jujube flesh offers an abundant source of raw material for flavonoid extraction, though it also contains other compounds such as organic acids. An optimized extraction process is essential to maximize the retention of active flavonoid components, providing a solid theoretical foundation for the broader utilization of non-medicinal parts of Ziziphus jujube. This study applied the Box–Behnken method, following single-factor experiments, to design and refine the extraction procedure. The optimal conditions identified were as follows: a 60% ethanol concentration, an SLR of 1:25 g·mL−1, and a 60 min extraction period, resulting in a maximum TFZJF yield of 1.98%. As the crude extract contained TFZJF, further enrichment and purification were required for subsequent pharmacodynamic evaluations, demonstrating the simplicity, precision, and practicality of the extraction method.

The ethanol reflux extraction of TFZJF has the advantages of good solubility, the separation of impurities to improve the purity of the target substance, a high extraction yield, and it can retain its related activity. In this study, the optimal extraction conditions were optimized based on the single-factor experiment using the Box–Behnken method. The best extraction conditions are as follows: a 60% ethanol concentration, SLR of 1:25 g·mL−1, and 60 min extraction time. The maximum yield of TFZJF is 1.98%. Since the extracted material obtained by optimizing the extraction process is a crude extract containing TFZJF, it needs to be purified and concentrated. After purification by D101 macroporous resin, the purity reached 58.32%, indicating that the purification process has higher purification efficiency and is more suitable for purifying TFZJF. This can be used for subsequent pharmacological activity research, indicating that the extraction process is simple, accurate, and feasible.

Insomnia, a prevalent clinical condition, significantly disrupts daily functioning. Despite extensive research, its pathogenesis remains unclear, with several hypotheses proposed to explain its development [30]. The leading theories include monoamine or its receptors, neuroendocrine, neuronal damage, and cellular molecular mechanism hypotheses, with the “neurotransmitter hypothesis” being widely accepted among researchers [31,32]. Contemporary studies indicate that neurotransmitter balance directly influences sleep quality, as neurotransmitters mediate communication between neurons and are critical to brain function, particularly during sleep. Their release and regulation are directly linked to sleep quality [33,34]. Furthermore, research has identified a strong association between insomnia and the dysregulation of monoamine and amino acid neurotransmitters within the central nervous system, with significant reductions in 5-HT, GABA, and BDNF levels observed in insomniac mice [35]. 5-HT, a neurotransmitter integral to central nervous system function, influences sleep, cognition, and various physiological processes. Reduced 5-HT levels are implicated in conditions such as sleep disorders [36]. GABA, an inhibitory neurotransmitter, regulates neuronal excitability, maintaining the balance of brain activity and promoting relaxation and sleep. Diminished GABA levels are strongly correlated with sleep disorders and related symptoms [37]. Investigating the role of these neurotransmitters in the brain is critical for understanding sleep regulation [38]. BDNF, a neurotrophic factor primarily active in the central nervous system, plays a fundamental role in neuroplasticity and mood regulation [39]. Alterations in BDNF expression are linked to the onset and progression of neurological conditions, making it essential for evaluating nervous system health and conditions such as insomnia and memory disorders [40,41]. Non-targeted metabolomics research on serum and brain biomarkers in insomnia model rats has identified phenylalanine and tryptophan metabolism as the most significantly impacted pathways associated with insomnia [42–44].

The therapeutic efficacy of Chinese medicine extends beyond symptom relief, targeting the underlying internal imbalances within the body. Metabolomics analysis reveals that PCPA significantly disrupts metabolites, including L-glutamate, L-tyrosine, and L-phenylalanine, in mouse serum. Notably, L-glutamate, a key biomarker in insomnia, plays a central role in glutamate metabolism, which is closely linked to the pathogenesis and progression of insomnia [45]. The present findings demonstrated that TFZJF exerted a substantial regulatory effect on the disruptions of metabolites such as L-glutamate, L-tyrosine, and L-phenylalanine, indicating a potential therapeutic benefit for insomnia. Additionally, glutamate, as a precursor to GABA, undergoes conversion into GABA through the action of glutamate decarboxylase. Glutamate and GABA metabolism may reflect the equilibrium between neural excitation and inhibition [46]. A significant reduction in GABA levels further supports this, aligning with previous research. Additionally, TFZJF is implicated in regulating metabolic pathways related to phenylalanine, tyrosine, and alanine metabolism. L-phenylalanine, an essential amino acid, is metabolized into tyrosine via phenylalanine hydroxylase, which is subsequently converted to dihydroxyphenylalanine by tyrosine hydroxylase. This process leads to the production of excitatory neurotransmitters such as norepinephrine and dopamine, both of which are associated with insomnia [47,48]. In summary, TFZJF may exert therapeutic effects on insomnia by modulating key metabolites, including L-glutamate, L-tyrosine, L-phenylalanine, and their related metabolic pathways. Additionally, this study has some limitations, mainly including a small sample size. In future studies, we will increase the sample size to further improve the reliability and accuracy of the research results and provide a certain basis for the resource development and utilization of non-medicinal parts of Ziziphus jujube.

## Author Contributions

Conceptualization, Y.Y. and Z.S.; methodology, J.L. and Y.Y.; software, B.L.; validation, Y.Y., Z.S.; formal analysis, Z.S.; resources, Y.Y., Z.S.; data curation, B.L.; writing—original draft preparation, J.L.; writing—review and editing, Y.Y.; visualization, Z.S..; funding acquisition, Y.Y., Z.S. All authors have read and agreed to the published version of the manuscript.

## Funding

This research was funded by Shaanxi Provincial Key Research and Development Program (2024CY-JJQ-41); Modern agricultural industrial technology system construction project (CARS-21); Doctoral Science Foundation of Shaanxi University of Chinese Medicine (306-17102032236); Major Science and Technology Project Plan of Xianyang City (2018k01-41).

## Conflicts of Interest

The authors declare no conflicts of interest.

## Abbreviations

(GABA): γ-aminobutyric acid
(5-HT): 5-hydroxytryptamine
(ANOVA): Analysis of Variance
(BDNF): brain-derived neurotrophic factor
(CV): coefficient of variation
(DZP): diazepam
(DEM): Differentially expressed metabolites
(PCPA): para-chlorophenylalanine
(SLR): solid-liquid ratio
(TFZJF): Ziziphus jujube flesh

## Notes

### Competing Interest Statement

The authors have declared no competing interest.

